# CD33-CAR iNKT cells overcome acute myeloid leukemia heterogeneity by activating CD8+ T cells and enhancing tumor immunogenicity

**DOI:** 10.64898/2026.11.25.768846

**Authors:** Sisi Wang, Guangyu Zhou, Hao Yan, Rebecca Richards, Astrid Beerlage, Carolin Wanzenried, Romina Marone, Alejandra Gomez-Cadena, Thomas Longval, Chiara Bernardi, Amandine Pradier, Benedetta Fiordi, Chaiyaporn Boonchalermvichian, Astrid Melotti, Yara Hajj Younes, Pragallabh Purwar, Elena Sotillo, Mylène Docquier, Yves Chalandon, Lukas Jeker, Camilla Jandus, Crystal Mackall, Robert S. Negrin, Federico Simonetta

## Abstract

Acute myeloid leukemia (AML) exhibits extensive inter- and intra-patient antigenic heterogeneity, representing a major obstacle for chimeric antigen receptor (CAR)-based therapies. Cellular platforms that combine potent direct cytotoxicity with the ability to engage endogenous immune responses may help overcome this limitation. Invariant natural killer T (iNKT) cells are particularly attractive for allogeneic immunotherapy because of their lack of alloreactivity and their capacity to promote cross-priming of conventional T cells. To exploit these properties in AML, we generated CD33-directed CAR iNKT cells and demonstrated direct antitumor activity against CD33-expressing AML cells both *in vitro* and *in vivo*. Using an *in vitro* mixed-tumor model composed of parental CD33-expressing and CD33-knockout MOLM-14 cells we show that CD33-CAR iNKT cells efficiently eliminated CD33-positive leukemia cells but did not directly control the growth of CD33-knockout cells. However, the addition of conventional T cells, particularly CD8⁺ T cells, together with CD33-CAR iNKT cells resulted in robust T-cell activation and subsequent control of CD33-knockout MOLM-14 cells. Importantly, CD33-CAR iNKT cells induced marked phenotypic changes in antigen-negative leukemia cells, including upregulation of HLA class I and class II molecules, suggesting increased immunogenicity. Finally, we recapitulated both the antitumor and immunomodulatory effects of CD33-CAR iNKT cells in primary bone marrow samples from patients with newly diagnosed AML. Collectively, these findings show that CD33-CAR iNKT cells can promote CD8⁺ T-cell immunity against AML and support a therapeutic strategy capable of extending antitumor activity beyond the targeted antigen, thereby helping to overcome antigenic heterogeneity.

## Introduction

Chimeric antigen receptor (CAR) T cells have revolutionized the treatment of B-cell malignancies and multiple myeloma, but their use in other hematological malignancies, namely in acute myeloid leukemia (AML), still faces major challenges (1). AML is characterized by remarkable phenotypic heterogeneity (2), both between patients and within individual patients, where multiple leukemic subpopulations and subclones often coexist, each potentially displaying distinct antigenic profiles. Several AML-associated targets have been investigated, including CD33 (3), CD123 (4), CLL-1 (5), CD70 (6), and CD7 (7). However, targeting a single antigen is unlikely to achieve durable disease control, as antigen-negative leukemic clones can escape immune pressure and drive relapse.

Invariant natural killer T (iNKT) cells represent a rare lymphocyte population, accounting for less than 1% of T cells in humans and mice. They express a semi-invariant T-cell receptor that recognizes glycolipid antigens presented by the non-polymorphic CD1d molecule. Because their antigen recognition is not HLA-restricted, iNKT cells exhibit no alloreactive potential and have been associated with protection from GvHD in preclinical models. Several groups have successfully engineered iNKT cells with CAR constructs targeting both hematologic and solid malignancies(8–14). CAR-iNKT display several advantages over conventional CAR T cells including their ability to target tumor cells through both the CAR and the endogenous invariant T cell receptor (TCR) (14) as well as through the activating receptor NKG2D (15). Importantly, we previously demonstrated that CD19 CAR-engineered iNKT cells possess a remarkable ability to induce endogenous host immune responses (16). Specifically, CD19 CAR-iNKT cells promote cross-activation and cross-priming of host cytotoxic CD8^+^ T cells, resulting in robust epitope spreading and the generation of long-lasting antitumor immunity that persists beyond the physical persistence of the CD19 CAR-iNKT cell product itself. These findings have subsequently been validated by an independent group in additional murine tumor models (17). Notably, both studies showed that allogeneic CAR-iNKT cells outperform conventional CAR T cells in their capacity to induce endogenous antitumor immunity *in vivo* in immunocompetent murine models of cancer (16,17). The ability to recruit and amplify host immune responses therefore appears to be a distinctive and particularly potent characteristic of CAR-iNKT cells.

Given their capacity to induce immunity against tumor antigens beyond the CAR-targeted antigen, CAR-iNKT cells might be especially effective in AML. We hypothesized that, by stimulating endogenous T-cell responses against multiple leukemia-associated antigens, CAR-iNKT cells could potentially overcome the limitations of single-antigen targeting strategies. To test this hypothesis, we generated CAR-iNKT cells targeting CD33, one of the most extensively studied AML antigens and a prototype target for myeloid-directed CAR therapies, and assessed their direct antileukemic activity against CD33-expressing AML cells as well as their ability to induce conventional T cell activity against CD33-negative clones.

## Materials and methods

### Cell lines

AML cell lines (MOLM-14, OCI-AML-2 and OCI-AML-3, a kind gift of Pr. Jerôme Tamburini) were cultured in RPMI 1640 media supplemented with fetal bovine serum (FBS) and 1% penicillin-streptomycin. MOLM-14, OCI-AML-2 and OCI-AML-3 cell lines were stably transduced with mCherry/luciferase reporter genes (Plasmid #75020, addgene). MOLM-14 cell line was stably transduced with GFP/luciferase reporter genes (Plasmid #75021, addgene). To generate MOLM-14 CD33 knockout (KO), MOLM-14 cells (1 × 10e6) were resuspended in 20 µl Lonza supplemented SE electroporation buffer with 2 µg BE mRNA and 100 pmol sgRNA (sgRNA_CD33_KO1 CCACUCACCUGCCCACAGCA) and electroporated using the 4D-Nucleofector system (Lonza) with program CL-120. Immediately after electroporation, the cells were incubated for 10 min at room temperature for recovery. Cells were then transferred in a 6 well plate containing 3 ml complete media.

### Human CD33 CAR-iNKT cell generation

Healthy donor peripheral blood was obtained at the Blood Transfusion Center, Geneva University Hospitals (HUG). Peripheral blood mononuclear cells were collected by the Ficoll-Paque density gradient centrifugation. iNKT cells were enriched by magnet sorting targeting the Vα24Jα18+ invariant chain using anti-Vα24J18+ biotin (13-5806-82, clone 6B11, eBioscience) and streptavidin beads (557812, BD Biosciences). CD14+ monocytes were enriched from negative fraction using anti-CD14 biotin and streptavidin beads. The positively selected iNKT cells was placed in culture with a 1:1 ratio of autologous monocytes prepared from the negative fraction. Cell culture was stimulated with 50 ng/ml anti-CD3 antibody (317347, Biolegend) and 50 ng/ml anti-CD28 (302934, Biolegend) and 150 IU/ml IL-15 (PeproTech) in RPMI1640 media supplemented with 10% FBS and 1% penicillin-streptomycin. After 48 hours, activated iNKT cells were transduced by culturing them for 48 hours in retronectin (T100B, Takara)-coated 24-well plates with retrovirus encoding second-generation anti-CD33-CD28-CD3ζ CAR with IL-2 (300 U/mL, PeproTech). On day 7 of iNKT cultures, autologous monocytes were loaded with 200 ng/ml α-Galactosylceramide (α-GalCer, MERCK) for 2-4 h, irradiated and added to the culture at a ratio of 1:1. Cells were split twice per week with fresh media containing 150 IU/ml IL-15 and cryopreserved until use. The purity of the iNKT cells was evaluated by flow cytometry using antibodies PE anti-TCR Vα24 (IM2283, Beckman coulter), FITC antiTCR Vβ11 (IM1586, Beckman coulter), APC-R700 anti-CD3 (565119, BD Biosciences), BV480 anti-CD4 (566104, BD Biosciences), PerCP/Cyanine5.5 anti-CD8 (344710, Biolegend). Dead cells were excluded from analysis using DRAQ7 dye (Thermo Fisher Scientific). Transduction efficacy of CAR expression on iNKT cells was evaluated by APC/PE-conjugated streptavidin (405207/405204, Biolegend) binding to biotinylated protein L (29997, Thermo Fisher Scientific). If purity was below 90%, iNKT cells were enriched by staining with biotinylated antiVα24Jα18+ followed by positive selection with streptavidin microbeads. Flow cytometry data were acquired using a BD LSRFortessa flow cytometer (BD Biosciences) and analyzed with FlowJo v10.10.0 (FlowJo LLC).

### Human T cell enrichment

For T cell enrichment, the cells were counted and enriched by the negative magnet sorting using anti-CD19 biotin (302204, Biolegend), antiCD14 biotin (367106, Biolegend) and streptavidin beads. The negative fraction was counted and cryopreserved for future use. For CD4 and CD8 subset enrichment, the cells were positively enriched by the magnet sorting using anti-CD4 biotin (344610, Biolegend) and anti-CD8α biotin (301004, Biolegend) respectively and streptavidin beads. The positive fraction was counted and cryopreserved for use. The purity of T cells, CD4 and CD8 subsets were determined by the flow cytometric staining using antibodies PerCP/Cyanine5.5 or APC-R700 anti-CD3 (560835/565119, BD Biosciences), BV480 anti-CD4 (566104, BD Biosciences) and PerCP/Cyanine5.5 anti-CD8 (344710, Biolegend). Dead cells were excluded from analysis using DRAQ7 dye (Thermo Fisher Scientific).

### *In vitro* tumor growth inhibition assay

For CD33 CAR-specific anti-tumor evaluation, mCherry-transduced AML cell lines were co-cultured with either fresh or cryopreserved CD33 CAR-iNKT cells or untransduced iNKT cell counterparts at different effector/target ratios (E:T) starting from 2:1. Non-treated (NT) tumor cells were seeded as a control. For antigen heterogeneity model, target population was a 1:1 mixture of MOLM-14-WT (GFP+) and MOLM-14 CD33KO (mCherry+) cells. The target population was co-cultured with CD33 CARiNKTs at a 1:1 E:T ratio, supplemented with third-party donor conventional T cells (or their CD4/CD8 compartment) at an 8:1 E:T ratio. Control groups included non-treated MOLM-14 cells (co-culture of GFP+ MOLM-14 and mCherry+ MOLM-14 CD33KO cells), MOLM-14 cells with conventional T cells (ratio 8:1) and MOLM-14 cells with CD33 CAR-iNKT cells (ratio 1:1). CD33 CAR-iNKT cell number was determined based on transduction efficacy. Conventional T cell or CD4/CD8 subtype number was calculated based on the CD3/CD4/CD8 purity. The realtime cell growth of all cell lines was measured using mCherry and/or GFP and the Incucyte S3 live-cell analysis instrument (Sartorius).

### *Ex vivo* AML primary blast killing assay and phenotypic analysis

Primary AML bone marrow samples were obtained from AML patients at initial diagnosis in Geneva University Hospitals (HUG) with written informed consent. Bone marrow mononuclear cells were collected by the Ficoll-Paque density gradient centrifugation and cryopreserved for future use. The primary AML patient samples were analyzed for blast cell phenotype using flow cytometry. Blast cells were identified as SSClow and CD45dim cells. Cells were stained with BUV737 anti-HLA-A2 (741869, BD Biosciences), BV421 anti-CD34 (343610, Biolegend), BV480 anti-CD45 (566115, BD Biosciences), BV605 anti-CD38 (562665, BD Biosciences), BV711 antiCD117 (313230, Biolegend), BV786 anti-CD3 (565491, BD Biosciences), FITC anti-CD33 (303303, Biolegend), PE anti-CD1d (561757, BD Biosciences), PE-Cy7 anti-HLA-DR (335830, BD Biosciences), APC anti-CD123 (567275, BD Biosciences). Dead cells were excluded from analysis using DRAQ7 dye (Thermo Fisher Scientific). For primary blast cytotoxicity, AML blasts and CD33 CAR-iNKT cells were co-cultured at 1:1 ratio using opposing HLA-A2 expression (either HLA-A2+ AML with HLA-A2-CAR-iNKT, or vice versa) as a phenotypic surrogate for specific population gating. At 48 hours, cells were harvested, stained with above antibodies, and the number of live tumor cells were determined using CountBright Absolute counting beads (C36950, invitrogen) by flow cytometry.

### *In vitro* MHC expression modulation

The 1:1 mixture of MOLM-14-WT (GFP+) and MOLM-14 CD33KO (mCherry+) cells were cocultured with CD33 CAR-iNKT cells at a ratio of 1:1. At day 7, cells were collected and stained with the antibody panel: APC-R700 anti-CD3 (565119, BD Biosciences), PE-Cy7 antiHLA-DR (335830, BD Biosciences), APC anti-HLA-ABC (55555, BD Biosciences). Dead cells were excluded from analysis using DRAQ7 dye (Thermo Fisher Scientific).

### *In vitro* T cell activation and proliferation assay

Cryopreserved conventional T cells were thawed and stained with CellTrace violet dye (C34557, invitrogen). T cells and CD33 CAR-iNKT cells with differential HLA-A2 expression were added into the antigenic heterogeneic tumor model with the same ratio as killing assay. At day 7, cells were collected and stained with the antibody panel: APC-R700 anti-CD3 (565119, BD Biosciences), BV480 anti-CD4 (566104, BD Biosciences), PerCP/Cyanine5.5 anti-CD8 (344710, Biolegend), APC anti-HLA-A2 (343308, Biolegend), BV605 anti-CD38 (562665, BD Biosciences), PE anti-ICOS (313507, Biolegend), PE-Cy7 anti-HLA-DR (335830, BD Biosciences). Dead cells were excluded from analysis using DRAQ7 dye (Thermo Fisher Scientific).

### RNA sequencing

MOLM-14 CD33KO cells were FACS-sorted at the end of the 7-days coculture. Total RNA was extracted from MOLM-14 CD33KO cells using the Direct-zol RNA Miniprep kit (R2050, Zymo Research) according to the manufacturer’s instructions. Sequencing libraries were generated using TruSeq Stranded mRNA Kit (Illumina) following the manufacturer’s protocol and sequenced on an Illumina platform to yield 100-bp single-end reads. Raw counts were compiled, retaining transcripts with at least 2 counts in at least one sample. The DESeq2 package was utilized to normalize data via variance stabilizing transformation (vst) for Principal Component Analysis (PCA) and to identify differentially expressed genes (DEGs) using a negative binomial generalized linear model. Significant DEGs, defined by an absolute log2 fold-change > 1 and a Benjamini-Hochberg adjusted P < 0.05, were visualized as volcano plots using the ggplot2 and ggrepel packages. To elucidate immunological signatures, Gene Set Enrichment Analysis (GSEA) was conducted. Genes were converted to Entrez IDs (clusterProfiler and org.Hs.eg.db), ranked by log2 fold-change, and interrogated against MSigDB Hallmark gene sets (msigdbr). Significantly enriched pathways (adjusted P < 0.05) and their transcript density distributions were visualized using ridgeplots via the enrichplot package.

### In vivo study

NSG mice (6-8 week old) were intravenously engrafted with mCherry+ MOLM-14 cell line 5x10^5^ cells per mouse. At day 3, fresh CD33 CAR-iNKT cells were injected through tail vein with 10x10^6^ cells per mouse adjusted to transduction efficacy. All animal procedures were conducted in strict accordance with the guidelines of the Swiss Federal Animal Protection Act and were formally approved by the Cantonal Veterinary Office of Geneva under the licence N°GE482. Animals were monitored according to the statements in the animal licence GE482 for clinical signs of distress, including body weight loss, lethargy, dehydratation or severe posture changes. Animals were humanely euthanized accordingly to the score sheet approved by cantonal authorities.

### Statistical analysis

Statistical analyses were performed using R v4.5.1 and R studio version 2026.01.2. Appropriate statistical methods employed are indicated in figure legends.

## Results

### Generation and antitumor effect of human CD33 CAR-iNKT cells against AML

We first established a protocol to generate human CD33 CAR iNKT cells from healthy donors’ peripheral blood. After isolation and activation, iNKT cells were transduced with a CD33 CAR construct using a retroviral system. Cells were then restimulated with α-GalCer and further expanded. This approach allowed for the generation of CD33 CAR-iNKT cells that were highly pure (Figure 1A) and that displayed a robust transduction efficiency (Figure 1B). CD33 CAR-iNKT cell expansion varied across donors in terms absolute cell numbers at the beginning or at the end of the culture (Figure 1C). Flow cytometry analysis showed that the CD33 CAR-iNKT product composition varied substantially among donors, with most products being predominantly composed of double-negative or CD4⁺ iNKT cells, whereas CD8⁺ iNKT cells generally represented a minor fraction of the final product (Figure 1D). We next evaluated the *in vitro* antitumor activity of CD33 CAR-iNKT cells against the AML cell line MOLM-14, which expresses high surface levels of CD33 (Supplementary Figure 1A). The growth of MOLM-14 stably transduced with a mCherry reporter gene cultured with CD33 CAR-iNKT cells at different effector:target (E:T) ratios was measured longitudinally. CD33 CAR-iNKT cells inhibited tumor growth of MOLM-14 at 1:1 ratio and displayed a dose dependent antitumor effect (Figure 1E-F). These findings were confirmed in additional AML models, including the OCI-AML-2 cells expressing high levels of CD33 (Supplementary Figure 1B) and OCI-AML3 cells expressing lower levels of CD33 (Supplementary Figure 1C). In both additional models, CD33 CAR-iNKT cells induced robust, dose-dependent tumor growth inhibition (Supplementary Figure 1D-G). We next assessed the antitumor efficacy of CD33 CAR-iNKT cells *in vivo*. NSG mice were engrafted with MOLM-14 cells and treated intravenously with CD33 CAR-iNKT cells three days later (Figure 1G). As shown in Figure 1H, treatment with CD33 CAR-iNKT cells significantly prolonged mice survival compared with untreated controls (Figure 1H). Collectively, these results demonstrate that peripheral blood-derived iNKT cells can be efficiently engineered to express a CD33-specific CAR and exert significant antitumor activity against AML both *in vitro* and *in vivo*.

**Figure 1.**
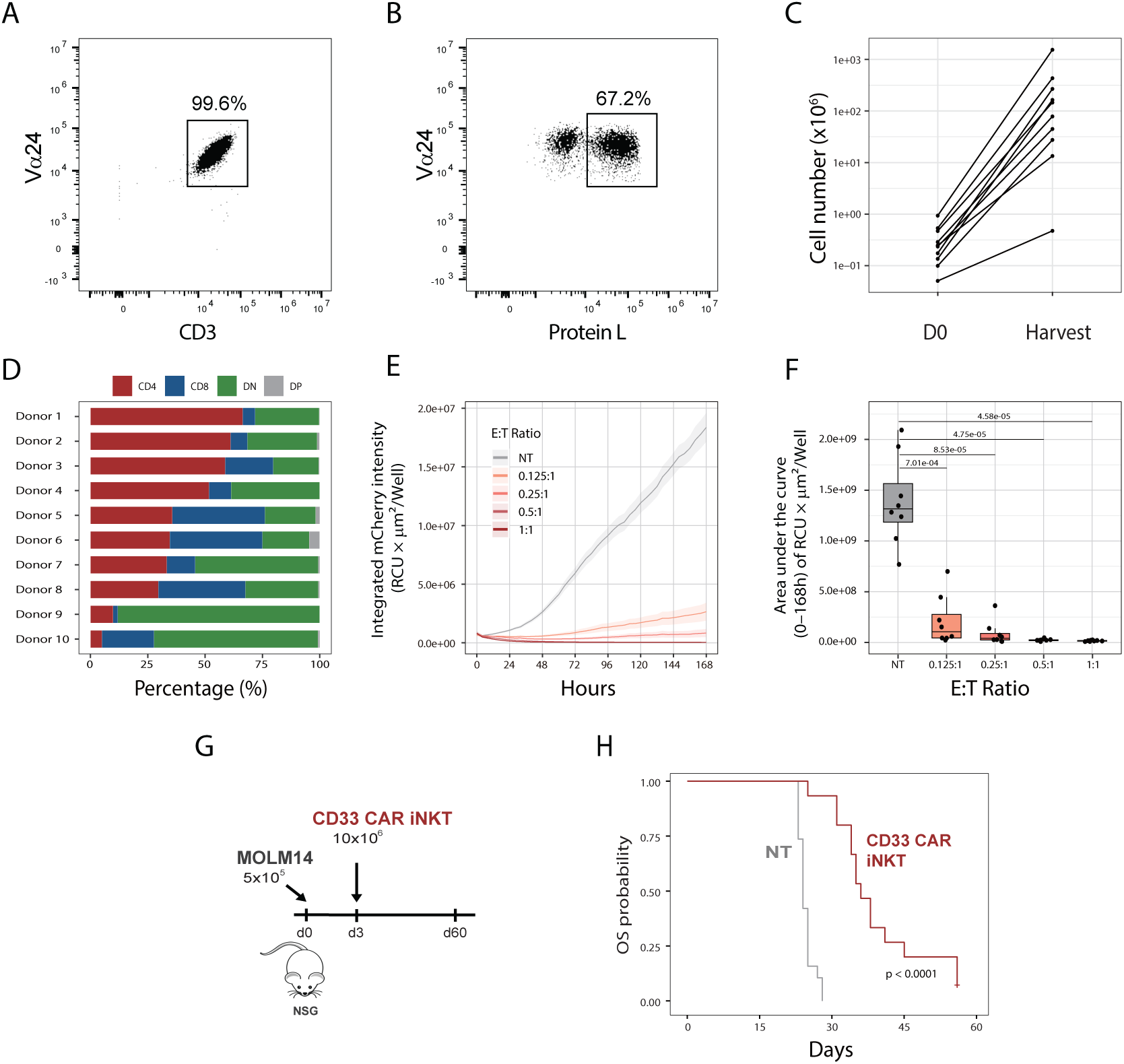
Generation and antitumor activity of human CD33-CAR iNKT cells. (A) Purity of expanded iNKT cells following ex vivo expansion, assessed by Vα24 and CD3 staining. (B) Transduction efficiency of CD33-CAR iNKT cells determined by Protein L staining. Representative flow cytometry plots from one of 10 healthy donors. (C) Absolute numbers of CD33-CAR iNKT cells at day 0 and at the end of expansion across different healthy donors. (D) Distribution of CD33-CAR iNKT cell subsets based on CD4 and CD8 expression. (E) *In vitro* growth of MOLM-14 cells cultured alone (NT) or with CD33-CAR iNKT cells at the indicated effector-to-target (E:T) ratios. (F) Tumor growth was quantified as the area under the curve (AUC) of integrated fluorescence intensity. Statistical comparisons were performed using the T test. Data are summarized from eight independent experiments. (G) Schematic representation of the *in vivo* experiment evaluating the antitumor activity of CD33-CAR iNKT cells. NSG mice were engrafted with MOLM-14 cells on day 0 and treated on day 3 with 10 × 10e6 CD33-CAR iNKT cells, normalized for transduction efficiency. (H) Survival of mice engrafted with MOLM-14 cells and treated with CD33-CAR iNKT cells (red) or left untreated (NT, gray line). Data are pooled from two independent experiments comprising 8–10 mice per group. Survival curves were generated using the Kaplan–Meier method and compared by the log-rank test. P values are indicated.

### Indirect antitumor effect of CD33 CAR-iNKT cells by engaging allogeneic conventional T cells

CAR iNKT cells have been reported to exert CAR-independent antitumor effects through NKG2D-dependent mechanisms (15). To assess the potential contribution of CAR-independent mechanisms in the effects of our cellular product, we compared the activity of CD33 CAR-iNKT cells against parental MOLM-14 cells or CD33-knockout MOLM-14 cells (Figure 2A). CD33 CAR-iNKT cells completely suppressed the expansion of CD33-expressing parental MOLM-14 cells but demonstrated no impact on tumor growth of CD33KO MOLM-14 (Figure 2B), confirming that the anti-tumor effect was CD33-dependent with negligible if any contribution of CAR-independent mechanisms. A critical advantage of the CAR-iNKT platform lies in its innate immunomodulatory capacity by recruiting endogenous T cell to exert prolonged tumor control (16).We hypothesized that by engaging T cells, the therapeutic impact of CD33 CAR-iNKT cells can extend beyond the primary target and circumvent AML antigen heterogeneity. To test our hypothesis, we established a mixed-culture system consisting of equal numbers of GFP-expressing parental MOLM-14 cells and mCherry-expressing CD33-knockout MOLM-14 cells. These tumor cells were cultured with CD33 CAR-iNKT cells in the presence or absence of third-party allogeneic conventional T cells (Figure 2C). As expected, CD33 CAR-iNKT cells efficiently controlled the growth of CD33-expressing parental GFP-MOLM-14 cells regardless of the presence of allogeneic conventional T cells (Figure 2D, upper panels). In contrast, neither CD33 CAR-iNKT cells alone nor allogeneic conventional T cells alone exerted meaningful antitumor activity against the antigen-negative CD33-knockout MOLM-14 cells (Figure 2D, lower panels). Interestingly, however, the combination of CD33 CAR-iNKT cells and allogeneic conventional T cells resulted in partial but significant control of CD33-knockout MOLM-14 cells (Figure 2D, lower panels), indicating that CD33 CAR-iNKT cells can cooperate with allogeneic conventional T cells to exert an antitumor activity against antigen-negative AML cells. Importantly, this indirect antitumor effect required engagement of the CAR with its cognate antigen. Indeed, when cultures contained only CD33-knockout MOLM-14 cells, neither CD33 CAR-iNKT cells alone nor their combination with conventional T cells were able to induce tumor control (Supplementary Figure 2A). Similarly, untransduced iNKT cells failed to promote antitumor activity against CD33-knockout cells even in the presence of allogeneic conventional T cells (Supplementary Figure 2B). Collectively, these data demonstrate that activation of CD33 CAR-iNKT cells through recognition of CD33-positive AML cells can induce bystander responses of allogeneic conventional T cells capable of targeting CD33-negative tumor cells.

**Figure 2.**
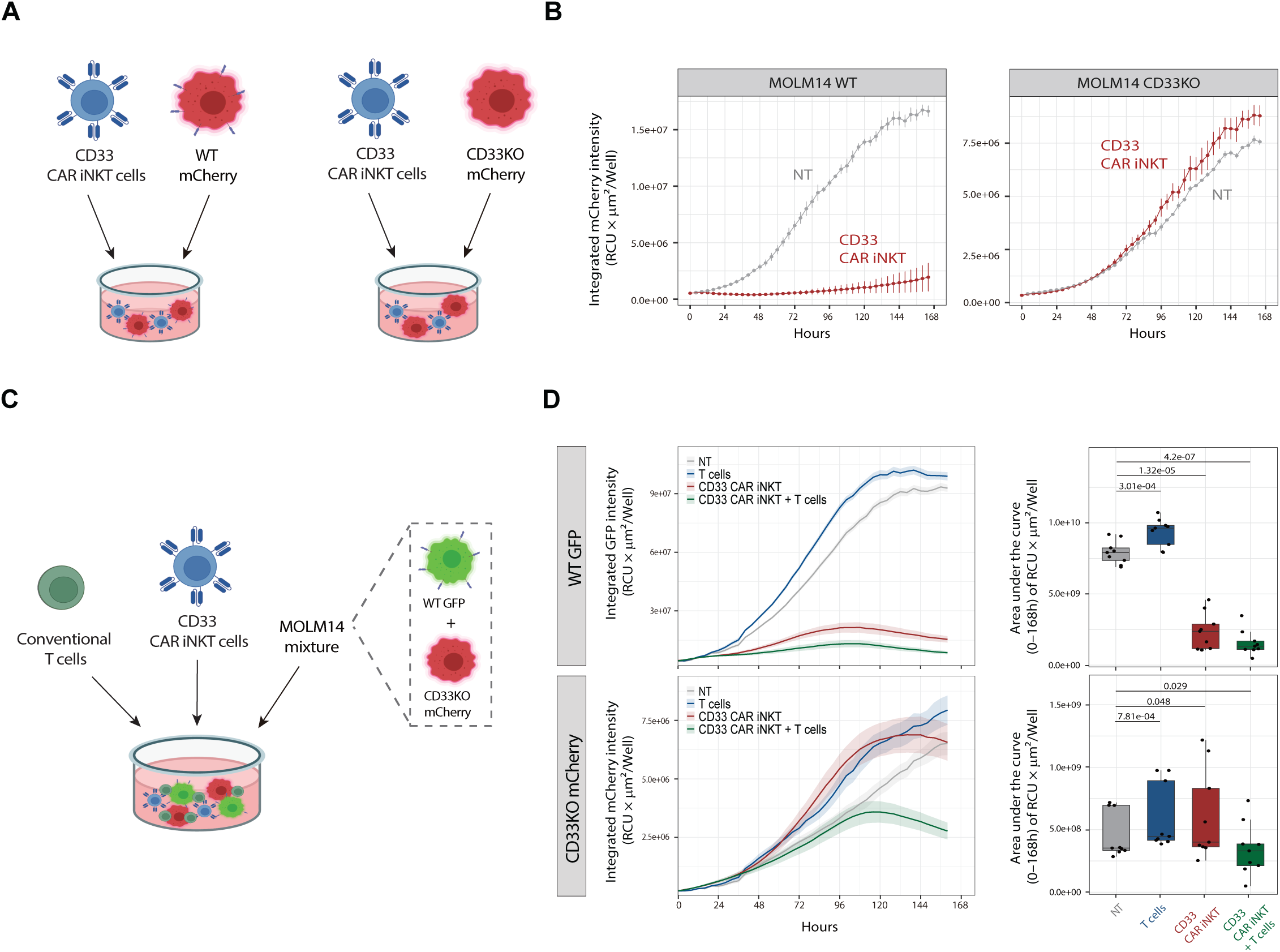
CD33-knockout AML cells escape direct CD33-CAR iNKT cell cytotoxicity but are controlled in the presence of conventional T cells. (A) Schematic representation of the experimental design. (B) Longitudinal growth of parental CD33-expressing MOLM-14 cells (left panel) or CD33-knockout MOLM-14 cells (right panel) cultured alone (NT, gray line) or with CD33-CAR iNKT cells (red line). Representative of two independent experiments. (C) Schematic representation of the mixed-culture model. GFP-expressing parental MOLM-14 cells and mCherry-expressing CD33-knockout MOLM-14 cells were co-cultured in the presence or absence of CD33-CAR iNKT cells and conventional T cells. (D) Longitudinal growth of parental GFP-MOLM-14 cells (upper panels) and CD33-knockout mCherry-MOLM-14 cells (lower panels) cultured alone (NT, gray line), with conventional T cells (blue line), CD33-CAR iNKT cells (red line), or both (green line). (E) Tumor growth was quantified as the area under the curve (AUC) of integrated fluorescence intensity. Data are summarized from nine independent experiments. Statistical comparisons were performed using the T test.

### CD33 CAR-iNKT cells promote activation of allogeneic conventional CD4⁺ and CD8⁺ T cells and enable CD8⁺ T cell-mediated control of antigen-negative AML cells

We next investigated the effects of CD33 CAR-iNKT cells on allogeneic conventional T-cells in our co-culture system. Phenotypic analyses by flow cytometry demonstrated robust activation of both CD4⁺ and CD8⁺ T cells following co-culture with CD33 CAR-iNKT cells in the presence of AML cells. Expression of the CD38 and HLA-DR activation markers was significantly increased in T cells cocultured with CD33 CAR iNKT cells compared with T cells cultured alone or with tumor cells (Figures 3A–D). Analysis of T-cell proliferation revealed a statistically significant but modest effect in the proportion of proliferating CD4⁺ T cells in the presence of CD33 CAR iNKT cells (Figure 3 and C). Conversely, the co-culture with CD33-CAR iNKT cells was associated with the appearance of a sizeable and significantly increased population of proliferating CD8⁺ T cells that was virtually absent in the other culture conditions (Figure 3B and D). To determine the relative contribution of CD4⁺ and CD8⁺ T cells to the indirect antitumor effects observed in our *in vitro* antigen-escape model, we repeated the mixed-culture experiments using isolated CD4⁺ or CD8⁺ T cells. Neither CD4⁺ nor CD8⁺ T cells alone displayed significant antitumor activity against parental or CD33-knockout MOLM-14 cells (Figure 3E - upper panel). The combination of CD33 CAR-iNKT cells and CD8⁺ T cells resulted in substantial and nearly complete control of antigen-negative AML cells, whereas no significant tumor control was observed in the presence of CD4⁺ T cells (Figure 3E - lower panel). Collectively, these findings indicate that CD33 CAR-iNKT cells induced the activation of both CD4⁺ and CD8⁺ T-cell populations but preferentially promote proliferation and functional cytotoxic responses from the CD8⁺ T cell compartment.

**Figure 3.**
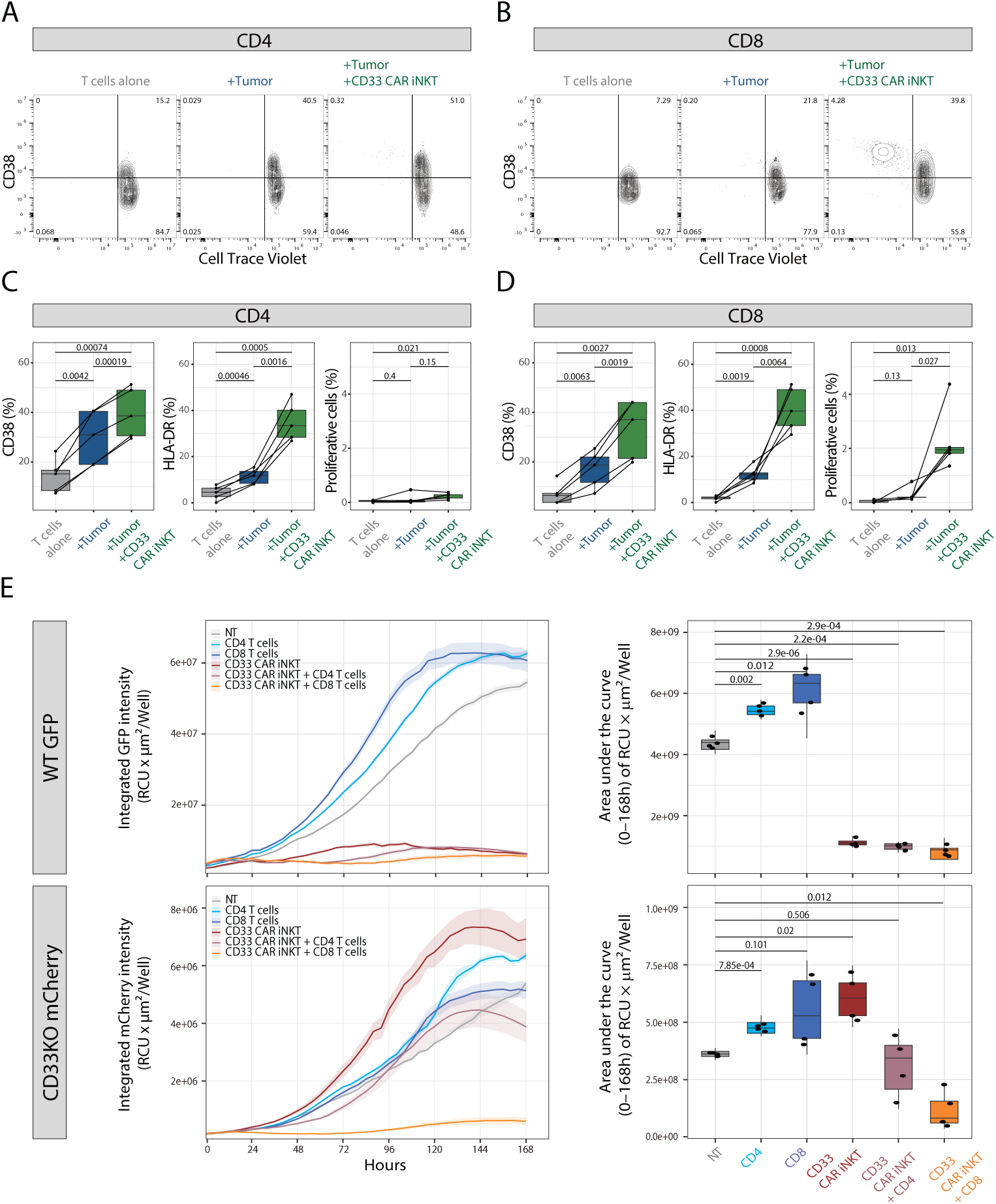
CD33-CAR iNKT cells activate CD8 T cells and promote elimination of antigen-negative AML cells. (A–B) Representative CellTrace Violet dilution and CD38 expression of CD4 (A) and CD8 (B) T cells cultured alone, with MOLM-14 cells, or with MOLM-14 cells and CD33-CAR iNKT cells. (C–D) Quantification of CD38 and HLA-DR expression and proliferation of CD4 (C) and CD8 (D) T cells. Data are summarized from five independent experiments. (E) Longitudinal growth of parental CD33-expressing GFP-MOLM-14 cells (upper panels) and CD33-knockout mCherry-MOLM-14 cells (lower panels) cultured alone (NT), with CD4 T cells, CD8 T cells, CD33-CAR iNKT cells, or the indicated combinations. (F) Tumor growth was quantified as the area under the curve (AUC) of integrated fluorescence intensity. Data are summarized from four independent experiments.

### CD33 CAR-iNKT cells modulate HLA expression on AML cells and promote allogeneic conventional T-cell responses

To better elucidate the mechanisms underlying the indirect antitumor activity mediated by CD33 CAR-iNKT cells, we next performed transcriptomic profiling on FACS-sorted CD33-knockout MOLM-14 cells recovered at the end of the seven days co-culture. Principal component analysis revealed a marked transcriptional impact of CD33 CAR-iNKT cell exposure on residual CD33-knockout AML cells (Figure 4A). While untreated MOLM-14 cells and MOLM-14 cells cultured with conventional T cells alone clustered closely together, cells exposed to CD33 CAR-iNKT cells displayed a distinct transcriptional profile, separating along both principal component 1 (49% of variance) and principal component 2 (23% of variance) (Figure 4A). Differential gene expression analysis performed between CD33KO MOLM-14 exposed to CD33 CAR iNKT cells and untreated MOLM-14 cells showed the upregulation of multiple genes involved in antigen presentation pathways, particularly genes encoding HLA class I and class II molecules (Figure 4B). To validate these observations at the protein level, we quantified surface expression of HLA molecules on MOLM-14 cells following co-culture with CD33 CAR-iNKT cells. Consistent with transcriptomic findings, exposure to CD33 CAR-iNKT cells resulted in significant upregulation of both HLA class I (HLA-ABC; Figure 4C) and HLA class II (HLA-DR; Figure 4D) molecules on residual MOLM-14 cells. Together, these results demonstrate that activated CD33 CAR-iNKT cells induce profound transcriptomic and phenotypic changes in AML cells by enhancing the expression of antigen-presentation molecules, potentially increasing their immunogenicity and their susceptibility to recognition by allogeneic conventional T cells. To further investigate the contribution of conventional T cells to the indirect effect of CD33 CAR iNKT cells, we compared the transcriptomic profiles of CD33-knockout MOLM-14 cells cultured with CD33 CAR-iNKT cells alone or in combination with allogeneic conventional T cells. Differential gene expression analysis revealed additional upregulation of multiple immune-related genes in the presence of allogeneic conventional T cells, including interferon-responsive genes and genes associated with inflammatory signaling pathways (Figure 4E). Gene set enrichment analysis identified significant enrichment of Hallmark pathways related to allograft rejection, interferon-γ response, interferon-α response, and oxidative phosphorylation (Figure 4F). Collectively, these findings indicate that activation of CD33 CAR-iNKT cells initiates a broader immune response enhancing the immunogenicity of CD33-negative AML cells and involving interferon-driven pathways induced by allogeneic conventional T cells.

**Figure 4.**
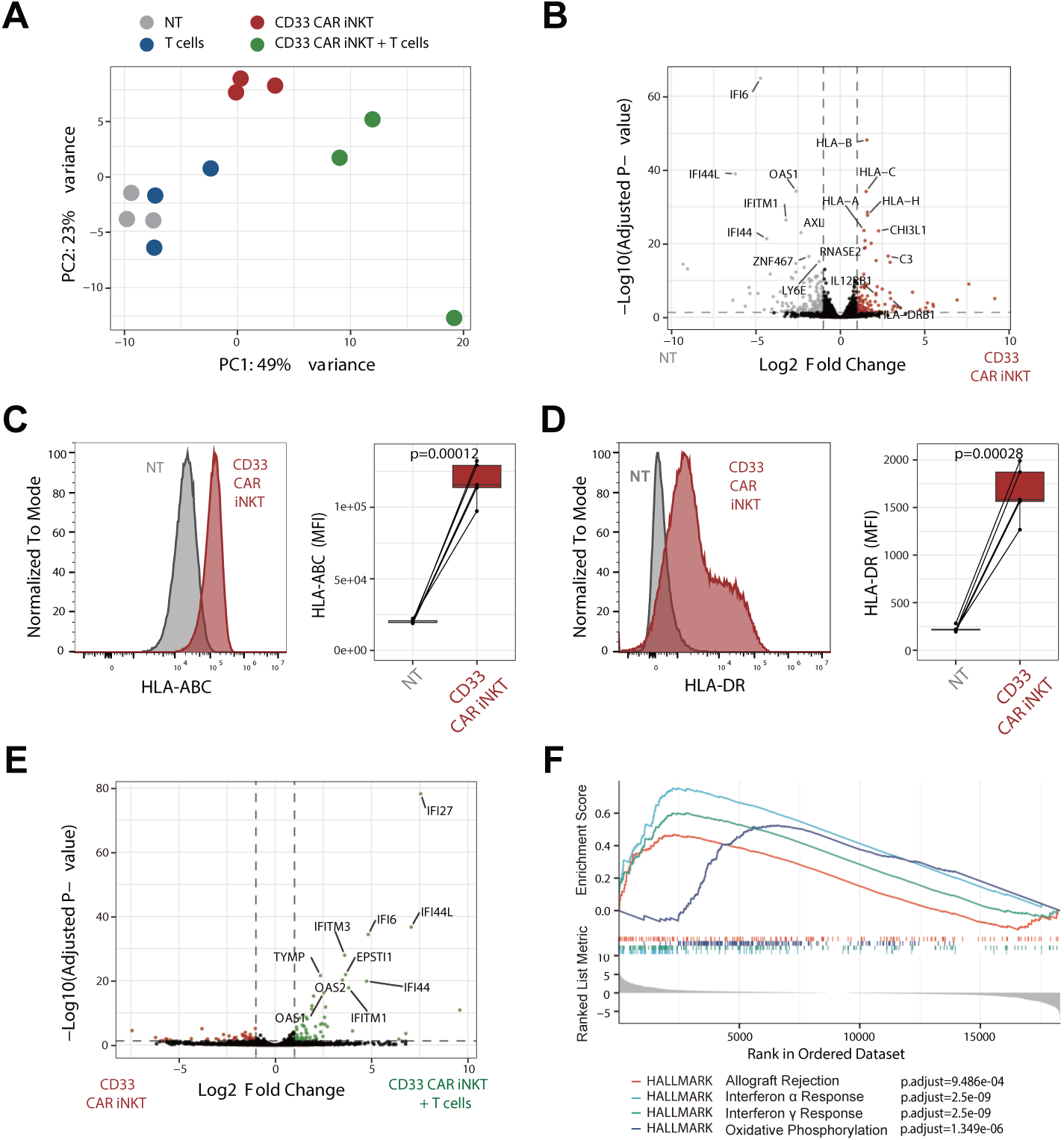
CD33-CAR iNKT cells induce an immunogenic phenotype in residual antigen-negative AML cells. (A) Principal component analysis of transcriptomic data from residual CD33-knockout MOLM-14 cells cultured alone (NT, gray dots), with conventional T cells (blue dots), CD33-CAR iNKT cells (red dots), or both (green dots). (B) Volcano plot showing differentially expressed genes in residual CD33-knockout MOLM-14 cells cultured alone (NT) or with CD33-CAR iNKT cells. Selected immune-related genes are highlighted. (C-D) HLA class I (C) and II (D) expression on residual CD33-knockout MOLM-14 cells cultured alone (NT, grey histograms and boxes) or with CD33-CAR iNKT cells (red histograms and boxes). Left panels, representative histograms; right, quantification from five independent experiments. (E) Volcano plot showing differentially expressed genes in residual CD33-knockout MOLM-14 cells cultured with CD33-CAR iNKT cells in the presence or absence of conventional T cells. (F) Enrichment plots from gene set enrichment analysis (GSEA) for the hallmark gene sets showing significant enrichment of immune-related pathways, including allograft rejection, interferon-α response, interferon-γ response, and oxidative phosphorylation, in residual CD33-knockout MOLM-14 cells cocultured with CD33-CAR iNKT cells and T cells compared with cells cultured with CD33-CAR iNKT cells alone. Gene signatures were obtained from the Molecular Signatures Database (MSigDB).

### CD33 CAR-iNKT cells exert potent antitumor activity against primary AML blasts and promote immune activation in patient samples

To validate our findings in a more clinically relevant system, we evaluated the activity of CD33 CAR-iNKT cells against AML blasts from bone marrow samples obtained from patients at AML diagnosis. Bone marrow mononuclear cells from five AML patients were cultured in the presence or absence of CD33 CAR-iNKT cells. Following co-culture, we assessed by flow cytometry direct antitumor activity as well as changes in tumor-cell phenotype and endogenous immune-cell activation (Figure 5A). CD33 expression on AML blasts was variable across patient samples, ranging from only partial/low to high levels (Figure 5B). CD33 CAR-iNKT cells induced substantial reductions in AML blast viability after 48 hours of co-culture, resulting in median reduction of approximately 50% of viable tumor cells, although the magnitude of response varied among patients (Figure 5C). Consistent with antigen-specific targeting, residual blasts displayed significantly reduced surface CD33 expression compared with untreated controls (Figures 5D–E), indicating preferential elimination of CD33-positive leukemic blasts. We next examined whether CD33 CAR-iNKT cells induced phenotypic alterations in residual AML cells similar to those observed in our *in vitro* cell-line model. Indeed, co-culture with CD33 CAR-iNKT cells significantly increased expression of HLA class I (HLA-A2; Figures 5F–G) and HLA class II (HLA-DR; Figures 5H–I) on residual AML blasts. Finally, we assessed the activation of endogenous T cells present within patient bone marrow samples upon exposure to CD33 CAR-iNKT cells. Coculture of bone marrow cells with CD33 CAR-iNKT cells resulted in significantly increased percentages of CD38+ and HLA-DR+ T cells compared with untreated controls (Figures 5J–L), demonstrating activation of endogenous T-cell populations. Taken together, these findings validate in primary patient samples the observations we made in our *in vitro* system and demonstrate that CD33 CAR-iNKT cells exert potent antitumor activity against CD33-expressing AML blasts while simultaneously enhancing tumor immunogenicity and promoting endogenous T-cell activation.

**Figure 5.**
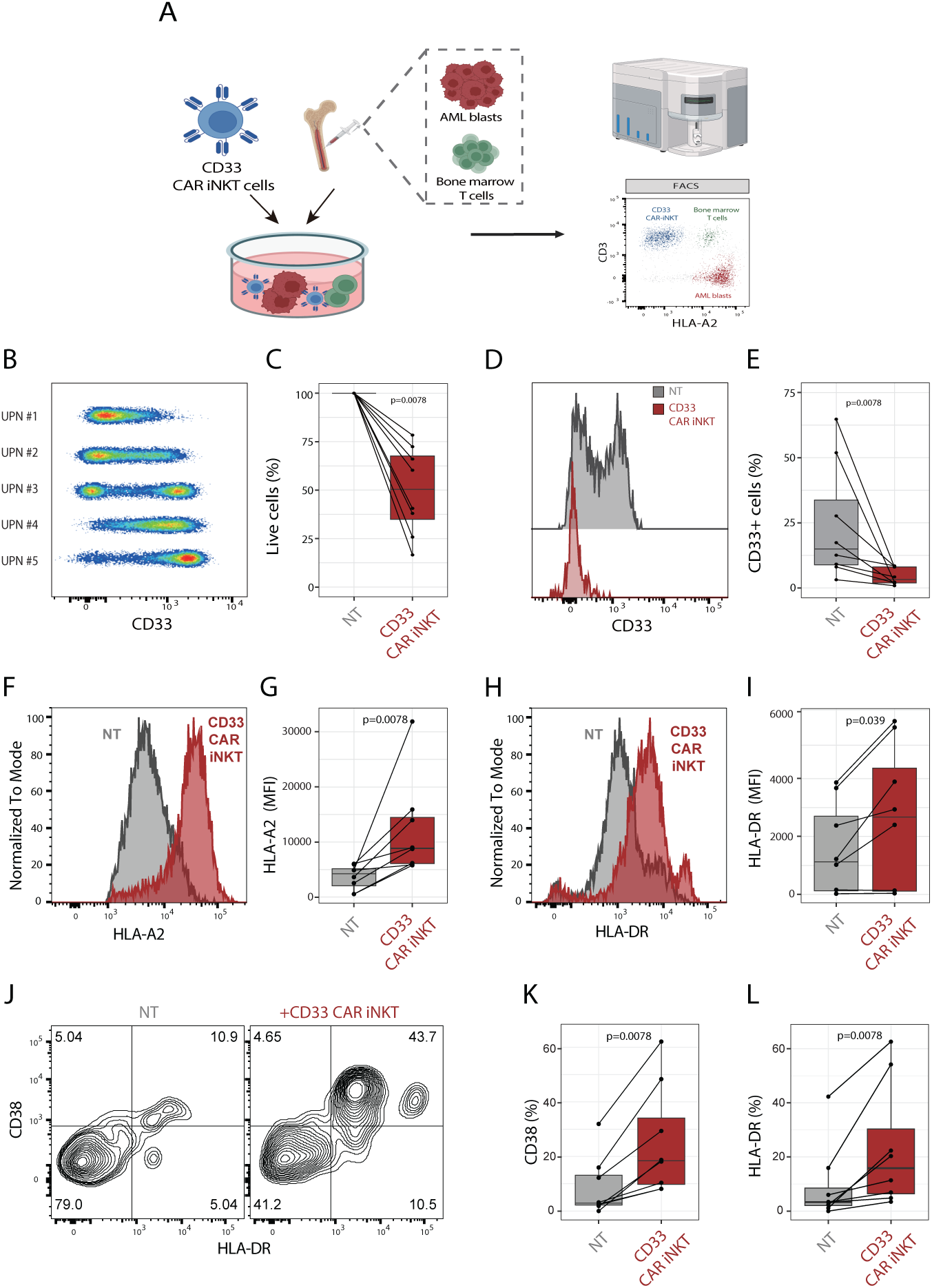
CD33-CAR iNKT cells exert antitumor and immunomodulatory activity in primary AML samples. (A) Schematic representation of the experimental design. CD33-CAR iNKT cells were co-cultured with freshly isolated bone marrow samples from patients with newly diagnosed AML. After 48 hours, flow cytometry was performed to quantify residual AML blasts and assess the phenotype of leukemic blasts and endogenous T cells. (B) Surface expression of CD33 on primary AML blasts from five patients. (C) Antitumor activity of CD33-CAR iNKT cells against primary AML blasts. Residual AML cells were quantified after 48 hours of coculture and data were normalized to NT controls. Data are summarized from eight independent experiments using samples from five patients and four different CD33-CAR iNKT cell products. (D) Representative histograms showing CD33 expression on residual AML blasts cultured alone (NT) or with CD33-CAR iNKT cells. (E) Quantification of CD33 surface expression on residual primary AML blasts. (F) Representative histograms showing HLA-A2 expression on primary AML blasts cultured alone (NT) or with CD33-CAR iNKT cells. (G) Quantification of HLA-A2 median fluorescence intensity. (H) Representative histograms showing HLA-DR expression on residual AML blasts cultured alone (NT) or with CD33-CAR iNKT cells. (I) Quantification of HLA-DR expression. (J) Representative plots showing CD38 and HLA-DR expression on endogenous T cells following co-culture with CD33-CAR iNKT cells. (K–L) Quantification of CD38 (K) and HLA-DR (L) expression on endogenous T cells. Data are summarized from eight independent experiments.

## Discussion

Our study demonstrates that CD33-CAR iNKT cells generated from peripheral blood-derived iNKT cells exert both direct and indirect antileukemic activity. In addition to efficiently eliminating CD33-expressing AML cells, CAR-iNKT cells promoted the activation of conventional T cells, resulting in immune control of antigen-negative leukemic cells. This indirect antitumor effect required CAR engagement by CD33-positive target cells and was mediated predominantly by CD8⁺ T cells.

Accumulating evidence indicates that the antitumor activity of CAR-iNKT cells extends beyond direct CAR-mediated cytotoxicity. Initial studies demonstrated that CAR-iNKT cells retain the ability to recognize CD1d-expressing tumor cells through their invariant T-cell receptor, thereby providing an additional mechanism of tumor recognition (14). More recently, independent groups have shown that CAR-iNKT cells can also mediate antitumor activity through NKG2D-dependent pathways, including modulation of NKG2D ligand expression on tumor cells (15,18). In our model, however, neither CAR-independent recognition nor NKG2D-mediated cytotoxicity could account for the elimination of CD33-negative AML cells, as CD33-CAR iNKT cells displayed no measurable cytotoxic activity against CD33-knockout MOLM-14 cells in the absence of conventional T cells. Instead, our findings identify recruitment and activation of endogenous T cells as the principal mechanism responsible for extending antitumor activity beyond the targeted antigen. These observations expand to a fully human AML model our previous findings, and those of others, demonstrating that CAR-iNKT cells can recruit bystander conventional T cells to participate in antitumor immunity (16,17).

We previously demonstrated in an immunocompetent murine lymphoma model that CAR-iNKT cells mediate both direct cytotoxicity and activation of endogenous immunity despite their relatively short *in vivo* persistence, allowing rapid recovery of normal antigen-expressing tissues (16). This feature may be particularly advantageous when targeting CD33, which is also expressed on normal myeloid progenitors. Unlike conventional CAR-T cells, whose prolonged persistence may result in sustained myeloablation, CD33-CAR iNKT cells may function as a transient immunotherapeutic bridge by providing immediate leukemia control while initiating endogenous immune responses capable of maintaining antitumor activity after disappearance of the cellular product. Such a strategy could preserve therapeutic efficacy while limiting long-term hematopoietic toxicity.

Previous studies have demonstrated the antileukemic activity of CAR-iNKT cells generated from human hematopoietic stem and progenitor cells (HSPCs), targeting antigens including CD33 (18) and CD70 (19). Our study complements these reports by demonstrating that peripheral blood-derived CAR-iNKT cells—the cellular source currently used in clinical development—possess potent immunomodulatory properties capable of engaging conventional T-cell responses. Whether HSPC-derived CAR-iNKT cells share these immunomodulatory functions remains unknown and warrants investigation before clinical translation.

Another notable finding was the ability of CD33-CAR iNKT cells to induce upregulation of both HLA class I and class II molecules on residual AML cells. Downregulation of HLA expression, particularly HLA class II, is now recognized as a major mechanism of immune escape following allogeneic hematopoietic stem cell transplantation (20,21). Restoration of HLA expression, potentially through interferon-γ and other inflammatory cytokines released during CAR-iNKT-mediated immune activation, may increase leukemia susceptibility to donor-derived T-cell responses and thereby enhance the efficacy of graft-versus-leukemia immunity of donor lymphocyte infusions. Although this hypothesis remains to be formally tested, it raises the intriguing possibility that CAR-iNKT therapy could synergize with post-transplant immunotherapeutic strategies.

Our study has several limitations. Although we reproducibly demonstrated activation of conventional T cells using primary human cells *in vitro*, validating this mechanism *in vivo* remains challenging. Xenograft models require profoundly immunodeficient hosts that lack endogenous adaptive immunity, whereas currently available humanized mouse models exhibit variable and often incomplete human immune reconstitution, including limitations in T-cell development and HLA-restricted immune responses, thereby limiting their ability to fully recapitulate human T-cell immunity (22). Consequently, our conclusions regarding induction of endogenous immunity rely primarily on robust *in vitro* observations, supported by complementary mechanistic evidence generated in immunocompetent murine models by our group and others (16,17).

Collectively, our findings demonstrate in a fully human experimental system that CD33-CAR iNKT cells can engage endogenous CD8⁺ T-cell responses capable of controlling leukemic cells independently of CAR target expression. By broadening antitumor immunity beyond direct CAR-mediated recognition, CAR-iNKT cells may overcome one of the principal mechanisms of resistance to CAR-based therapies, namely antigen-negative escape. More broadly, these results highlight the potential of CAR-iNKT cells to function not only as cytotoxic effector cells but also as initiators of endogenous adaptive immunity. This dual mode of action provides a strong rationale for the clinical development of allogeneic CD33-CAR iNKT cell therapy for AML, particularly in settings where antigen heterogeneity and immune escape limit the efficacy of conventional CAR-based approaches.

---

**Supplementary Figure 1.**
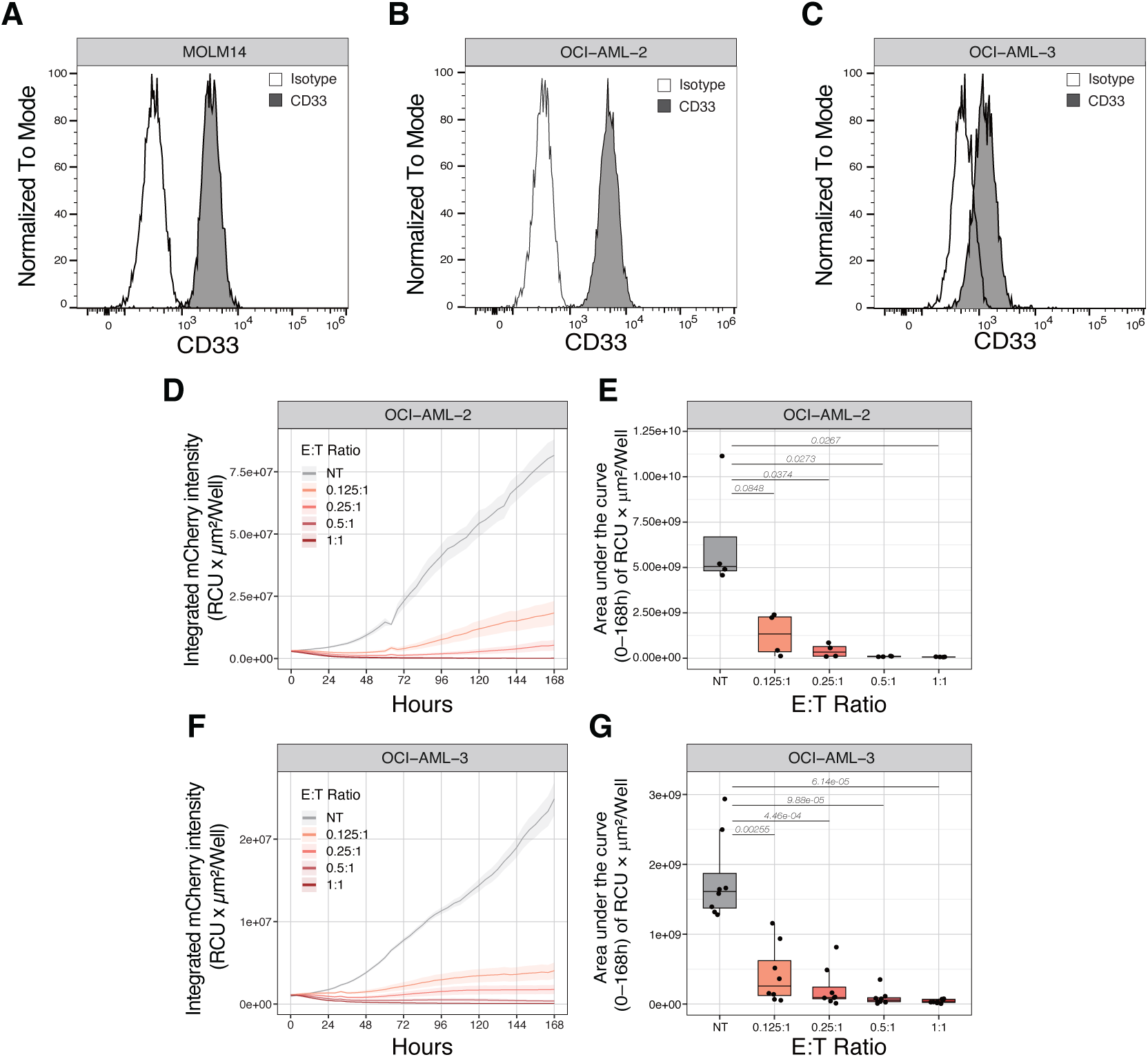
CD33 expression and antitumor activity of CD33-CAR iNKT cells in additional AML models. (A–C) Representative histograms showing CD33 expression on MOLM-14 (A), OCI-AML-2 (B), and OCI-AML-3 cells (C). (D–E) Antitumor activity of CD33-CAR iNKT cells against OCI-AML-2 cells. Left, representative longitudinal growth curves; right, tumor growth quantified as the area under the curve (AUC) of integrated fluorescence intensity. Data are summarized from four independent experiments. (F–G) Antitumor activity of CD33-CAR iNKT cells against OCI-AML-3 cells. Left, representative longitudinal growth curves; right, tumor growth quantified as the area under the curve (AUC) of integrated fluorescence intensity. Data are summarized from eight independent experiments.

**Supplementary Figure 2.**
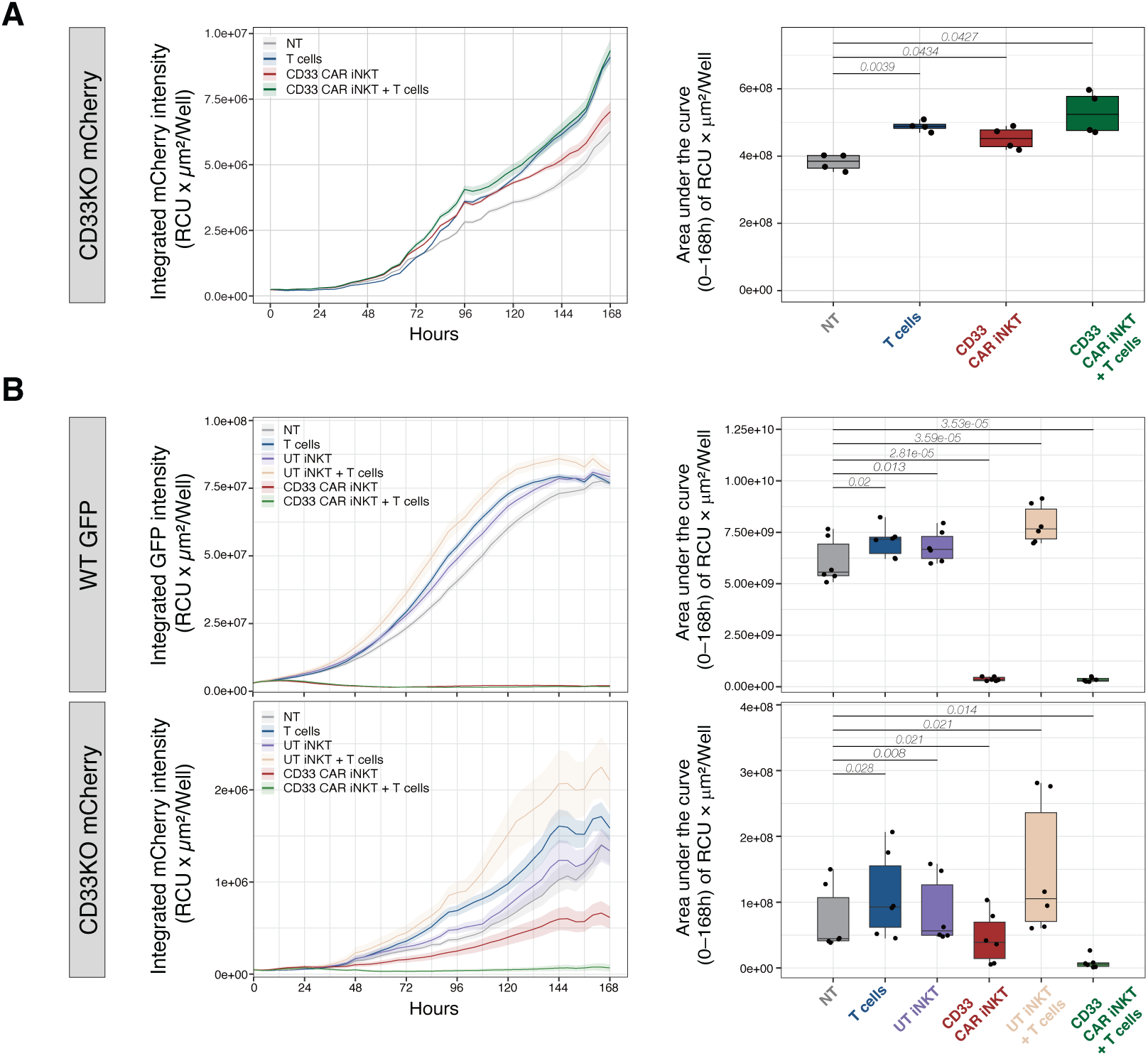
CD33-CAR engagement is required for activation of conventional T cells. (A) Longitudinal growth of CD33-knockout MOLM-14 cells cultured alone (NT), with conventional T cells, CD33-CAR iNKT cells, or both. (B) Comparison of the effects of untransduced and CD33-CAR iNKT cells on parental CD33-positive and CD33-knockout MOLM-14 cells in the presence or absence of conventional T cells. Left, representative longitudinal growth curves; right, tumor growth quantified as the area under the curve (AUC) of integrated fluorescence intensity. Data are summarized from six independent experiments.

## Financial support

This study was supported by the Fondation Dr Henri Dubois-Ferrière Dinu Lipatti to FS; Fond’Action to FS; SAKK/BMS Grant «HEM Pioneer» Award to FS; Novartis Foundation for Medical-Biological Research to FS; NIH PPG CA 49605 to RSN; Fondation Henriette Meyer to SW and China Scholarship Council to SW.

## Notes

### Competing Interest Statement

The authors have declared no competing interest.

